# Sequence-dependent Stability and the Apparent Two-state Thermal Transition of Extended Collagen Triple Helices

**DOI:** 10.64898/2026.07.29.741348

**Authors:** Sophie Youngji Xu, Sam Wong, Sally Tan, Fahmida Akter, Yujia Xu

## Abstract

The thermal stability of collagen triple helices is strongly influenced by the amino acid sequence of the repeating Gly–X–Y tripeptides, yet how these residue-specific interactions are integrated within an extended triple helix to determine thermal behavior remains poorly understood. Here, we addressed this question using recombinant collagen mimetic peptides (rCMPs) containing extended native sequences from the α_1_(I) and α_2_(I) chains of human type I collagen. Triple-helix formation was nucleated by a C-terminal foldon domain and further stabilized by interchain disulfide crosslinking, allowing the apparent melting temperature (*Tₘ*) to reflect interactions within the triple-helical domain independent of nucleation. The stabilizing effects of Pro and Y-position Arg identified in host-guest peptides were largely preserved in extended triple helices, whereas the proposed Lys–Gly–Glu (KGE) interchain salt bridge produced little measurable stabilization, demonstrating the influence of sequence context. Remarkably, identical triple-helical sequences exhibited markedly different thermal behavior when unfolding was initiated under different conditions. Nevertheless, extended triple helices differing substantially in sequence and length retained an apparently two-state thermal transition. These findings support a mechanism in which unfolding is preferentially initiated within regions of lower intrinsic stability, while the continuity of the triple helix couples neighboring regions into a cooperative unfolding process throughout the helix. This mechanism provides a plausible explanation for the longstanding paradox that extended collagen triple helices exhibit persistent sequence-dependent thermodynamic heterogeneity despite a two-state thermal transition, and a framework for investigating how sequence-dependent stability contributes to the structure and function of collagen molecules.

**TOC:** 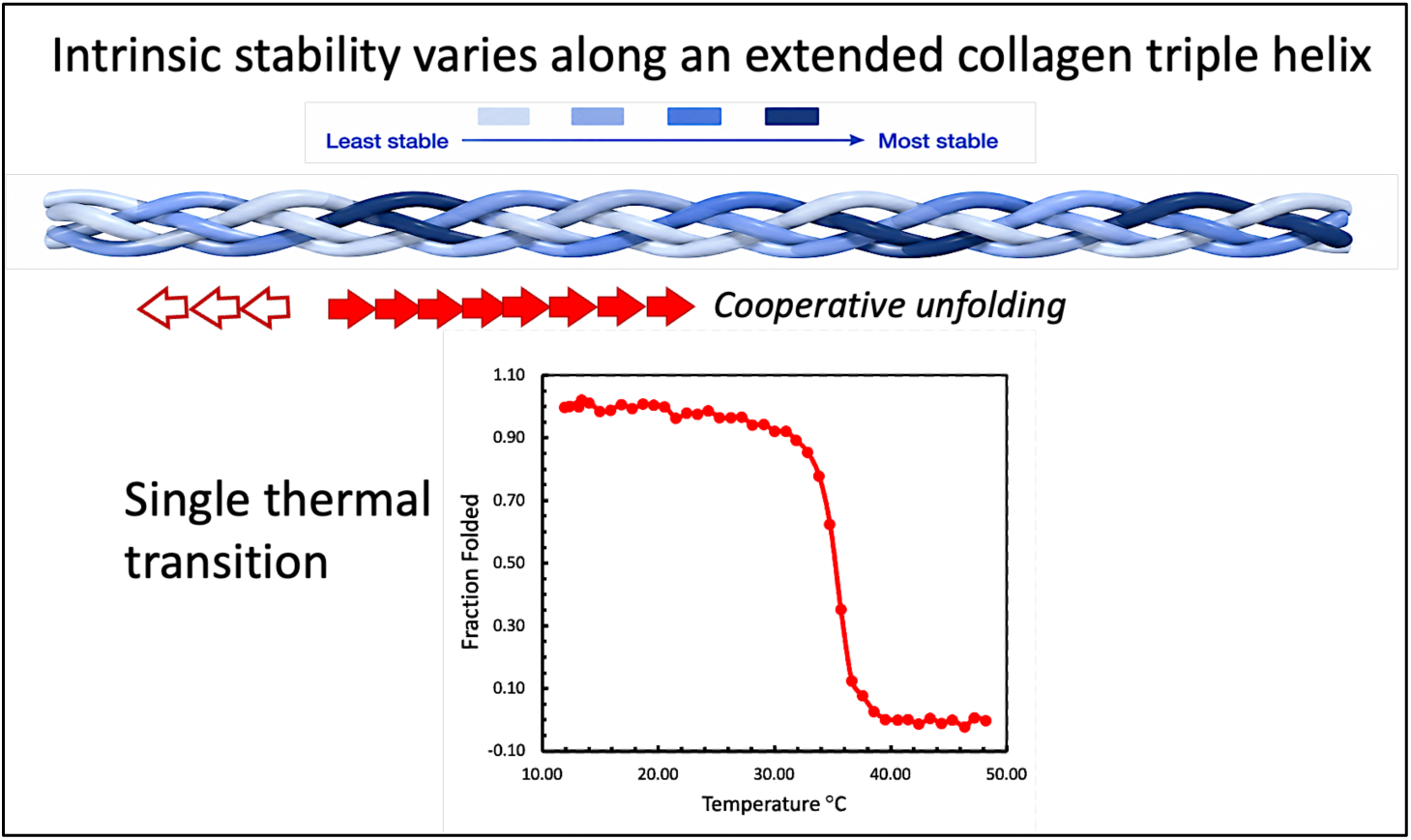

## INTRODUCTION

How residue-specific interactions are integrated to determine the stability of an extended collagen triple helix remains a central unresolved question in collagen biophysics. The concept that collagen triple helices contain regions of differing intrinsic stability has become central to understanding collagen function, disease, and biomaterial design(1–7). Yet despite its broad acceptance, experimental identification and characterization of such domains has remained surprisingly elusive. Thermal denaturation of native fibrillar collagens, even those with highly heterogeneous amino acid sequences, typically exhibits a single cooperative transition that closely resembles two-state unfolding, obscuring any underlying structural heterogeneity (6, 8). This apparent paradox arises from the unique architecture of the collagen triple helix. Unlike most proteins, collagen consists of three parallel polypeptide chains wrapped around a common axis with a tightly packed glycine core, a nearly uniform backbone geometry, and few long-range tertiary interactions. This structural simplicity imposes a strict sequence requirement—Gly must appear at every third residue within uninterrupted Gly–Xxx–Yyy (GXY) repeats, and native collagens often consist of over 300 GXY repeats within each polypeptide chain. Triple-helix stability is maintained by a network of main-chain hydrogen bonds, whereas residues at the X and Y positions modulate stability through interchain side-chain interactions and by restricting the polyproline II (PPII)-like backbone conformation (9–12). The central challenge is to understand how cooperative stability domains emerge within a continuous collagen triple helix and how their local unfolding is translated into the global thermal transition observed experimentally. Understanding how these sequence-dependent interactions are manifested in the cooperative thermal behavior of extended collagen triple helices has therefore remained a major experimental challenge.

Most current knowledge of sequence-dependent stability derives from chemically synthesized, short collagen-model peptides (CMPs) stabilized by flanking repeating (Gly–Pro–Hyp) sequences, which generally restrict native sequence insertions to only 6–9 residues(13, 14). Within these simplified systems, residue contributions often appear approximately additive (15, 16). Such observations naturally support a view in which triple-helix stability is determined by the sum of local residue interactions. While this framework has been highly successful in defining residue-specific stability and for guiding the design of a wide variety of CMPs, it provides little basis for understanding how sequence-dependent stability might become partitioned into distinct regions within longer native triple helices. Indeed, attempts to extrapolate these residue-specific measurements to extended collagen triple helices have met with limited success, leaving unresolved how local sequence effects are integrated withing long, native-like triple helices.

Here, we examine this question using protein-engineered recombinant collagen mimetic peptides (rCMPs) consisting of 63 residue Col-domains corresponding to residues 877–939 of the α_1_(I) and α_2_(I) chains of human type I collagen. By extending the accessible sequence context well beyond the ∼6–9 residue window of conventional CMPs, these rCMPs permit residue-specific effects to be evaluated within a substantially more native structural environment while preserving a continuous triple-helical segment in which the cumulative influence of multiple native sequences can be examined. Under conditions in which triple-helix conformation is nucleated at the C-termini, thereby minimizing contributions from nucleation itself and more closely resembling the unfolding of a fully folded native collagen triple-helix, we found that the stabilizing effects of certain individual residues remain robust and significant, whereas the contribution of other interactions, particularly interchain salt bridges, are more limited than suggested by short-peptide models and are dependent on the sequence context. More intriguingly, the pronounced susceptibility of the triple-helix to terminal destabilization suggests a model in which unfolding initiated within a thermolabile region propagates cooperatively along the helix, potentially explaining why native collagens predicted to contain heterogeneous stability regions nevertheless often exhibit an apparently two-state like thermal transition. Together, these results refine current model of collagen sequence-stability and provide a framework for understanding collagen function and for the rational design of native-like collagen mimetics and collagen-based biomaterials.

## MATERIAL AND METHODS

### Buffer preparation and peptide solutions

All peptides were first dissolved in 5 mM acetic acid, pH 3.6 to prevent aggregation and stored at ∼5 °C. For temperature melt experiments at different pH, the peptide solution was mixed with equal volume of 2X concentrated buffer, and the pH after mixing was confirmed by reproducible mixing test of equal volume of 5 mM acetic acid and corresponding buffers. The 2X phosphate buffer (PB) consists of 58 mM sodium dibasic phosphate and 42 mM sodium monobasic phosphate with no added salt; pH 7.0. The 2X phosphate saline buffer (PBS) consists of 58 mM sodium phosphate dibasic, 42 mM sodium phosphate monobasic, 254 mM NaCl and 54 mM KCl, pH 7.0. The 2X pH 9.4 phosphate buffer was made similar as the PB except the pH was first adjusted to 10.20 using NaOH; 2X pH 9.4 phosphate saline buffer was made the same way with the addition of salt (254 mM NaCl and 54 mM KCl). A separate mixing test confirmed the final pH is 9.4 after mixing with peptides in 5 mM acetic acid.

### Expression and purification of rCMPs

#### The expression plasmid

the codon optimized genes of the peptides were synthesized (GenScript) and inserted between BamHI and Ncol sites downstream from the T7 promoter site of pET32a(+) plasmid. The in-frame insertion of the gene was confirmed by DNA sequencing (GenWiz).

#### Expression

A single colony of transformed cells were inoculated into 10 ml LB+Amp media (containing 50 mg/ml ampicillin) for 18 hours with 225 RPM shaking at 37 ℃, and then transferred into 1L of LB+Amp media and grown in a shaker at 37 ℃ until the optical density at 600 nm reached 0.6-0.8 AU. To induce the expression,100 uL of 1M IPTG (isopropyl-β-D-1-thiogalactopyranoside) is added to the 1 L growth media. The media was then transferred to a shaker-incubator set at 16 ℃, 225 RPM shaking for 20 hours.

#### Purification

The cells were harvested by centrifugation at 2000 RCF, 4 ℃ for 20 minutes. The cells were resuspended in 10 mL lysis buffer (50 mM Tris with 300 mM NaCl), with the addition of 100 uL of 100 mM PMSF (phenylmethylsulfonyl fluoride), 1 mL of 20 mg/mL lysozyme, 20 uL of 1mg/mL DNAse, and sonicate for complete lysis (Vibracell, 45 second of pulse with 20 seconds of break, repeated three times with machine set up of output 3.0, Duty cycle 50%). The lysed cells were centrifuged at 7800 RPM at 4 ℃, for 50 minutes to separate the cell debris from the soluble proteins. The supernatant was collected in a 50 mL centrifuge tube, and 2 mL Nickel-NTA resin slurry (ThermoFisher Scientific) was added and incubated on a shaker for 3 hours at 4 ℃ allowing the His-tagged protein to bind to the resin. The resin was washed twice with 10 mL lysis buffer, followed by 10 mL of 10 mM imidazole and then 10 mL of 50 mM of imidazole to remove the unspecific binding. The His-tagged protein was eluted using 10 mL of 500 mM of imidazole and collected in 1mL fractions. The elutions were analyzed using SDS_PAGE as described below. Fractions with high concentrations of protein were pooled and added to a dialysis cassette with 2800 Da molecular weight cutoff (Thermo Scientific™ Slide-A-Lyzer™ Dialysis Cassettes) and dialyzed in 4 L of 5 mM acetic acid for two hours at room temperature. The buffer was changed twice and after the second change, the whole setup was placed at 4 °C for 24 hours to completely remove imidazole.

After dialysis, the concentration was measured using Thermo Scientific NanoDrop OneC Microvolume UV-Vis Spectrophotometer and equilibrated at 4 °C for approximately two weeks. The molecular weight and the length of each peptide is summarized in Table S1.

#### High-performance Liquid Chromatography (HPLC)

Following the affinity column, peptides showing minor impurities were further purified using HPLC. The polar solvent used in HPLC is distilled water with 0.1% trifluoracetic acid (TFA), and the non-polar solvent used is acetonitrile. Total of 5 mL of protein sample was injected into a C-18 column and eluted at a 2 mL per minute rate. The peptides were eluted at ∼ 43% acetonitrile with a retention time of 23-27 minutes monitored by OD at 280 nm. The elutions were pooled, the purity confirmed by SDS PAGE gel electrophoresis, and lyophilized. The lyophilized powder was dissolved in 5 mM acetic acid at a concentration of ∼ 1 mg/mL and equilibrated at 4 °C for ∼ two weeks before use.

### SDS-PAGE

The purity of the proteins is confirmed by SDS-PAGE gel electrophoresis (Fig. S1). Samples were prepared by mixing 50 uL of protein samples with 12.5 uL of 5X DSD loading buffer and heated on the heat block set at 100 °C for 30 minutes. When reducing SDS buffer were used, the 5X SDS buffer contained 0.5M DTT. For the electrophoresis, 8 uL of samples were loaded onto a 0.75 mm thick 15% gel. Broad Multi Color Pre-Stained Protein Standard from Genescript was used as the molecular weight standard. The gels were run at constant voltage of 180 V and stained with Coomassie gel staining solution (0.1% Coomassie Blue R-25 in 45% methanol and 10% glacial acetic acid) for 30 minutes and destained in destaining solution (10% methanol, and 10% glacial acetic acid) for 1 hour.

### Temperature melting experiments

Circular dichroism spectroscopy was used to study the structure and the thermal stability of the triple helix formation (Chirascan V100). The CD-spectra were taken from 190 nm to 260 nm using 1 mm quartz cuvettes at 10 °C, with a scan rate of 1 nm per step per second (heating rate of 1 degree/min). The thermal denaturation of the triple helix monitored using temperature step-ramp mode with the settings of 1 °C per step, and 60 seconds equilibration time at each temperature. The CD signal (in millidegree) was normalized to mean residue ellipticity (*MRE*) using the following equation:

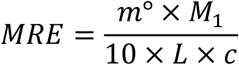

Where the *m*^°^ is CD signal in millidegree, *M_1_* is the molar mass of a single peptide chain, *c* is the concentration of protein in mg/mL, and *L* is the path length of the quartz cuvettes in mm. The concentration used were within the range of ∼ 0.2 mg/mL to 0.4 mg/mL.

Temperature melt experiment: The thermal transition curve is converted to fraction folded (FF) using the following equation and the apparent melting point (*T_m_*) was determined as the temperature (*T*) when the *FF* signal drops to 50%:

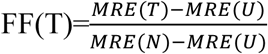

where *MRE(T)* is the signal of the sample at *T*, MRE(U) and MRE(N) are the MRE of the native and unfolded states at *T* based on the linear extrapolation from the native and unfolded baseline, respectively. The native baseline was obtained by a linear fit of the data between ∼ 5 °C-20 °C, while that of the unfolded state was the linear fit between ∼ 40 °C-50 °C.

The thermal transitions, which are independent of peptide concentration because of the foldon domain and the Cys-knot cross linking, were repeated 3-5 times for each peptide with samples from different preparations and in different concentrations (∼ 0.1–0.4 mg/mL). Typical reproducibility of the melting curves is shown in Fig. S2.

## RESULTS AND DISCUSSION

### The sequence-dependence of triple helix thermal stability

We developed two recombinant collagen mimetic peptides (rCMPs), α1C and α2C, each containing a 63-residue Col-domain modeling the section of residues 877–939 of the α_1_(I) and α_2_(I) chains of human type I collagen, respectively (Fig. 1A). Both peptides share the same overall design: a C-terminal foldon domain for nucleation and folding, a C-terminal Cys-knot (Gly–Pro–Cys–Cys) for interchain crosslinking (17), a (Gly–Pro–Pro)₄ (GPP₄) and a (Gly–Pro– Pro)_3_ (GPP_3_) segment flanking, respectively, the N- and C-termini of the Col-domain, and an N-terminal (His)₆ tag. Each peptide is 133 residues in length, with an 84-residue (i.e. 28 GXY repeats) triple-helical region comprising the Col-domain and the GPP₄ and GPP_3_ segments. This region of type I collagen contains numerous *Osteogenesis Imperfecta* (OI) mutations in which similar Gly substitutions can lead to markedly different clinical outcomes, suggesting an important role for local sequence context in modulating conformational dynamics (18, 19). The segment contains only two potential Hyp residues, both within the α_2_(I) chain, permitting expression in bacterial systems lacking proline hydroxylation.

**Figure 1:**
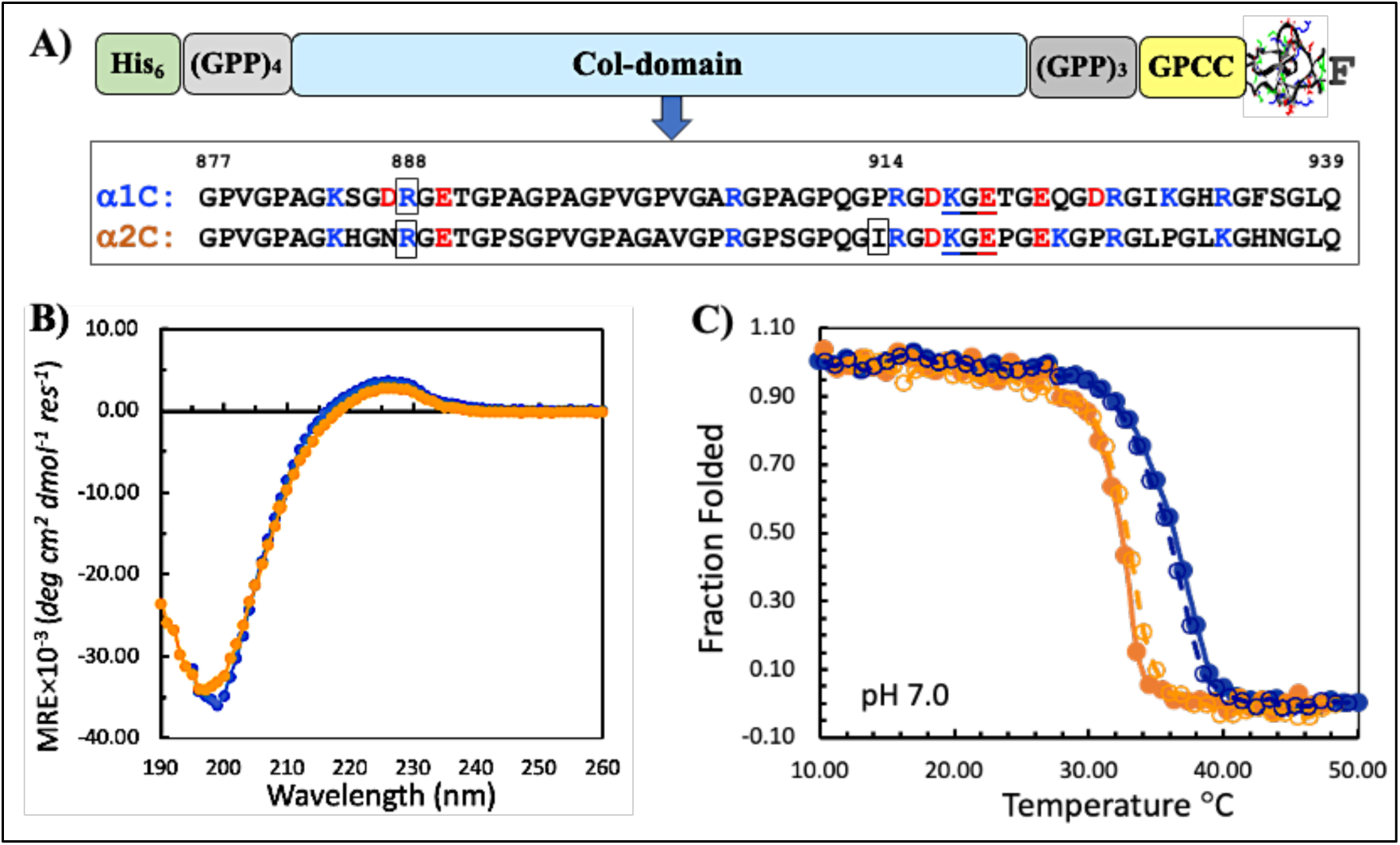
rCMP α1C and α2C. A) A schematic drawing of the primary structure of α1C and α2C; the amino acid sequences of the Col-domain are shown in the box with the positively and negatively charged residues highlighted in blue and red, respectively; the residues that are later mutated are boxed with their positions within the triple helix domain of human α_1_(I) shown above. The foldon domain (F) is shown as the stick and ribbon presentation created using SPDBV (PDB ID: 1RFO) with the backbone in grey, positively and negatively charged residues in blue and red, respectively, and polar residues in green. GPCC is the sequence inserted to form Cys-knot in the triple helix; GPP_4_ and GPP_3_ are (Gly-Pro-Pro)_4_ and (Gly-Pro-Pro)_3_ sequences, respectively, included to increase the stability of the triple helix. B) the CD spectra of α1C (blue) and α2C (orange) at 4 °C in 50meM phosphate buffer at pH 7.0. C) the temperature melting curves of α1C (blue) and α2C (orange) monitored by the change of ellipticity at 222 nm. The solid lines and filled symbols are in pH 7.0 phosphate buffer, the dashed lines and open symbols are in pH 7.0 PBS buffer which contains 127 mM NaCl and 27 mM KCl.

Both peptides form stable triple helices, as indicated by nearly identical CD spectra with a shallow positive peak at 225 nm and a strong negative peak at 197 nm (Fig. 2A), consistent with a well-folded triple helix with low Hyp content. Despite this structural similarity, their thermal stabilities differ markedly. The apparent melting temperature (*Tₘ*) of α1C is 37.36 ± 0.22 °C, whereas that of α2C is 33.57 ± 0.40 °C under identical buffer conditions (Fig. 1C). The addition of 155 mM of salt did not have noticeable effects on the thermal stability (Fig. 1C). This ∼4 °C difference in *T_m_* is notable given that 18 of the 28 GXY triplets are identical and sequence homology at the X and Y positions exceeds 80% (Fig. 1A).

**Figure 2:**
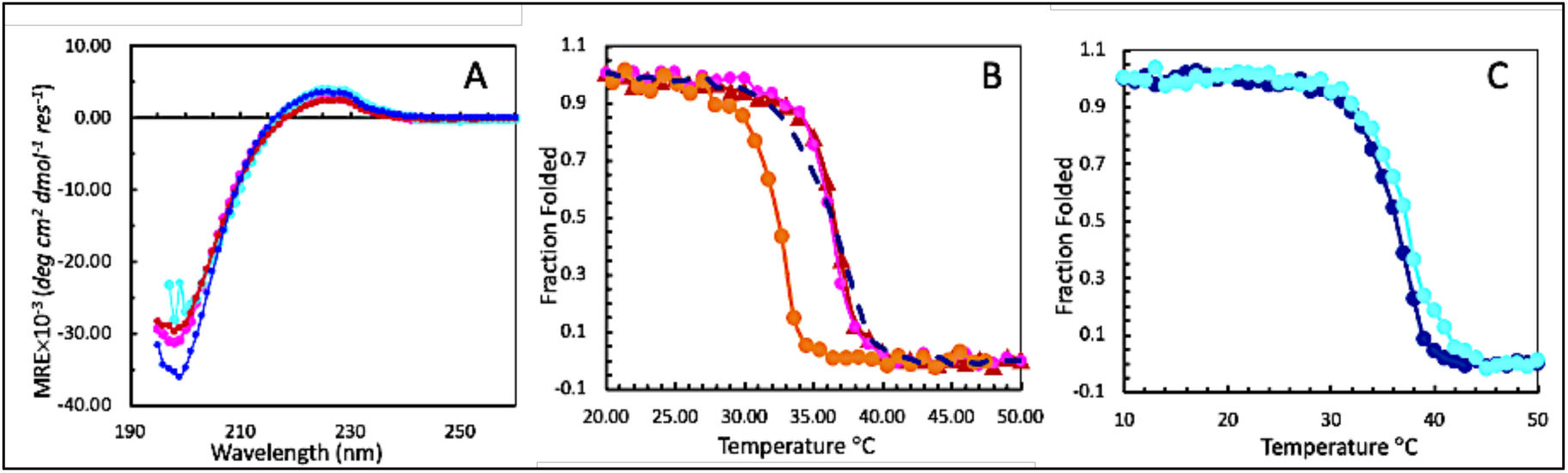
the muted effects of KGE sequences on the thermal stability of triple helices. A) the CD spectra of α1C (blue), α1C-K(cyan), α2C-P (red), and α2C-PK (magenta) at 4 °C. B) the temperature melting curve of α2C (orange filled circle), α2C-P (red triangle), and α2C-PK (magenta filled circle), and α1C (blue dash line). D) the temperature melting curves of α1C (blue) and α1C-K(cyan). All samples are in pH 7.0 phosphate buffer no added salt.

This variation in *T_m_* highlights the sensitivity of triple-helix stability to the identity of residues at the X and Y positions. Since the peptides are nucleated at the C-termini by foldon and the Cys-knot, the thermal transitions are independent of peptide concentration and the measured *T_m_* primarily monitors the unfolding–refolding equilibrium of the triple-helix domain (18). The intrinsic “triple-helix propensity” of amino acids including Hyp at X- and Y-positions has been established using host–guest peptides (15). Structurally, the two positions are not equivalent: Y residues are more solvent-exposed, whereas X residues are packed closer to the helix core and more likely to participate in interchain interactions (20). In short synthetic peptides, the contributions of individual GXY triplets are approximately additive (21). This additivity, however, does not extend to longer sequences—the same rules predict unphysical *T_ₘ_* values (∼ −100 °C) for native collagens as well as for α1C and α2C peptides. We therefore focus on relative effects, comparing each GXY triplet in α1C to its counterpart in α2C at the same position. The apparent *Tₘ* value of a GXY triplet measured in host–guest peptides cannot be directly translated to a quantitative impact on the *T_m_* of a long triple helix; these values are used only as a qualitative measure of the triple helix propensity of a triplet. For example, tripeptide Gly–Pro–Ala at positions 892-894 and 907-909 (the numbering referring to position in the triple helix domain of human type I collagen, which was used to be consistent with previous studies) contributes more strongly to stability than its α2C counterpart Gly–Pro–Ser at the equivalent positions (Fig. 1A) indicated by a ∼5.9 °C higher *Tₘ* in host–guest peptides. Gly–Pro–Arg at location 913-915 appears to be the triplet in α1C with the greatest stability advantage over its α2C counterpart Gly–Ile–Pro (GIP) as demonstrated by its ∼10.6 °C higher *T_m_* in host-guest peptides. In contrast, some triplets in α1C are relatively destabilizing compared to α2C, such as Gly–Asp–Arg (position 886-888) and Gly–Ala–Arg (position 904-906) when compared to Gly– Pro–Arg in those positions in α2C. Taken together, these position-by-position differences yield only a modest net advantage: across the Col-domain, α1C is more stable than α2C by ∼1.7 °C. Additional factors which are not captured by the host-guest peptides are clearly involved.

To test how individual triplets contribute within an extended native sequence, we replaced the Ile residue of the GIP triplet at position 914 (Ile914) of α2C with Pro to generate peptide α2C-P (Fig. 1A); the tripeptide GPP has a *T_m_* ∼ 9 °C higher than a GIP in host-guest peptides. Peptide α2C-P formed a stable triple helix with a *T_ₘ_* of 37.25 °C, nearly identical to α1C and ∼3 °C higher than α2C (Fig. 2A and B, Table 1). This marked increase of thermal stability by the introduction of only one Pro, highlighting the stabilizing effects of imino acids and also suggested that the intrinsic propensities measured in short peptides can be, at least, qualitatively informative even within long native-like sequences. In this case, introduction of a single Pro residue was almost sufficient to offset the combined destabilizing effects of multiple factors in α2C.

**Table 1.** The *apparent* melting temperature of rCMP triple helices in °C.

| Peptide | pH 7.2 |  | pH 9.4 |  | pH 3.6 |  | pH 2.9 |  |
| --- | --- | --- | --- | --- | --- | --- | --- | --- |
| | $T_m$ | $\Delta T_m^*$ | $T_m$ | $\Delta T_m^*$ | $T_m$ | $\Delta T_m^*$ | $T_m$ | $\Delta T_m^*$ |
| $\alpha 1C$ | $37.36 \pm 0.22$ | - | $37.15 \pm 0.21$ | - | $31.75 \pm 0.50$ | - | $31.25 \pm 0.35$ | - |
| $\alpha 2C$ | $33.57 \pm 0.40$ | -3.79 | $34.14 \pm 0.32$ | -3.01 | $28.83 \pm 0.15$ | -2.92 | $23.83 \pm 0.29$ | -7.42 |
| $\alpha 2P$ | $36.25 \pm 0.40$ | -1.11 | $36.60 \pm 0.36$ | -0.55 | $30.37 \pm 0.26$ | -1.38 | $27.03 \pm 0.54$ | -4.22 |
| $\alpha 2P-K$ | $36.15 \pm 0.21$ | -1.23 | $35.88 \pm 0.18$ | -1.27 | $30.80 \pm 0.30$ | -0.95 | $25.00 \pm 0.00^{\#}$ | -6.28 |
| $\alpha 1C-K$ | $37.58 \pm 0.30$ | +0.22 | $37.00 \pm 0.35$ | -0.15 | $30.78 \pm 0.33$ | -0.97 | $29.25 \pm 0.00^{\#}$ | -2.00 |
| $\alpha 2N$ | $36.03 \pm 0.21$ | -1.33 | $35.95 \pm 0.10$ | -1.20 | $35.15 \pm 0.17$ | +3.45 | $35.00 \pm 0.50$ | +3.75 |
| $\alpha 1C-GPP_6$ | $41.08 \pm 0.38$ | +3.28 | $41.00 \pm 0.25$ | +9.25 | $37.75 \pm 0.00^{\#}$ | +6.00 | | |

<sup>#</sup>: the thermal melt experiments were conducted twice with identical estimated apparent $T_m$ values. Examples of reproducible melting curves are shown in Fig. S2.

Next, we examined another interaction known to strongly stabilize synthetic collagen peptides. By replacing Arg888 in α2C-P with Lys, we introduced a Lys-Gly-Glu (KGE) sequence to generate peptide α2C-PK (Fig. 1A). In host-guest peptides, KGE motifs can form interchain salt bridges that increase *T_m_* by as much as ∼15.4°C and have been widely used to engineer heterotrimeric collagen mimetic peptides (12). Surprisingly, however, α2C-PK showed a *T_m_* nearly identical to α2C-P (Fig. 2B, Table 1), indicating the absence of a measurable net stabilizing effects of KGE in this sequence context. We observed the same result when introducing a KGE sequence in the equivalent position in α1C: peptide α1C-K displayed essentially the same thermal stability as α1C (Fig. 2A and C, Table 1).

The absence of a measurable stabilizing effect from the KGE sequence suggests that these electrostatic interactions are highly context dependent. In synthetic peptides, KGE-mediated stabilization is typically strongest in regions rich in imino acids, particularly Pro and Hyp, which constrain backbone geometry and may optimize interchain salt-bridge formation (9–11). In contrast, the present peptides contain relatively few imino acids and no Hyp. This altered sequence environment may affect the local helical geometry and weakening the spatial alignment required for efficient interchain electrostatic interactions (Bella 2010). To create the new KGE sequence in α1C-K and α2C-PK, the Arg-to-Lys substitutions also replaced intrinsically more favorable triplets with less stabilizing ones (GDK versus GDR in α1C and GNK versus GNR in α2C-P) based on host–guest studies (15). It is not clear whether such one-residue substitutions would produce the behavior observed in Host-guest peptides as the Ile-to-Pro mutation did, one cannot rule out the possibility that modest stabilization from partial salt-bridge formation may be offset by the intrinsic destabilization associated with the altered triplets. Our previous study of a peptide containing the same Col-domain as α1C demonstrated the stabilizing effects of the KGE in position 918-920 when a Gly-substituting mutation was introduced at position 913, suggesting the function of at least partially formed salt-bridges in this sequence context (18). Together, these results indicate that both electrostatic stabilization and local triplet propensities are strongly influenced by sequence context in long native-like collagen helices.

It is important to note that the thermal stability measured using the rCMPs directly reflects the interactions among the three polypeptide chains that stabilize the triple helix without requiring a separate nucleation step. The foldon domain remains stably folded under the conditions used for the thermal melt experiments (22). The Cys-knot further covalently links the three chins through a set of interchain disulfide bonds, thereby maintaining chain association at the C-terminus even when the N-terminal regions being largely disordered (18). In contrast, the stability of synthetic collagen peptides lacking cross-linking is strongly influenced by the nucleation reaction, which itself depends on amino acid sequence. Unfolding of these rCMPs triple helices with a fully folded foldon-domain at the C-terminus is likely initiated from the N-terminus and propagated through a zipper-like process according to the nucleation-propagation model of helix folding. Nevertheless, we cannot rule out the possibility that the folding and unfolding of isolated segments of the long helix proceed bidirectionally during the transition (23, 24).

### Unfolding initiation and propagation

The thermal stability of these peptides showed a surprisingly high sensitivity to unfolding at the N-terminus. While there was little change in *T_m_* of all peptides between pH 7.0 and pH 9.4 (Fig. 1 and Table 1), with or without added salt, all peptides showed a marked ∼ 5-6 °C decrease in *T_m_* at pH 3.6 (Fig. 3A, Table 1) when the residues of His-tag are protonated. It was known from studies of synthetic peptides that positive charges at the N-terminus of the triple helix were particularly destabilizing (25). What is striking here is that the electrostatic repulsion generated by the His-tag is sufficient to overcome the contribution of the adjacent GPP_4_ sequence inserted to stabilize the triple-helix (Fig. 1A). To determine whether this loss of stability originated from unfolding initiated at the N-terminus, we created peptide α2N by moving the crosslinking GPCC sequence to the N-terminus, thereby generating an N-terminal Cys-knot adjacent to the His-Tag (Fig. 1A). As expected, α2N showed only minimal decrease of *T_m_* for about 0.5 °C at pH 3.6 (Fig. 3B and C, Table 1, more discussion in the next section).

**Figure 3.**
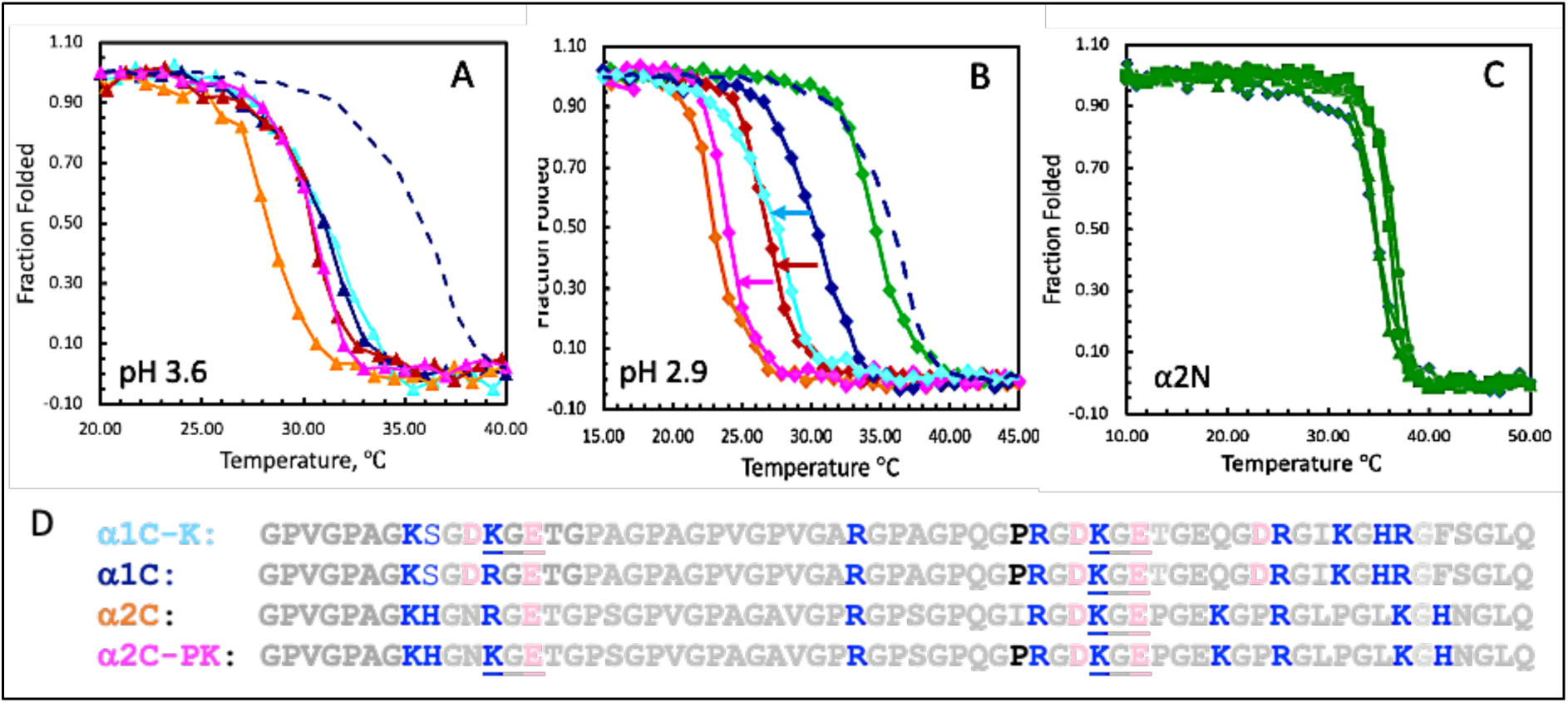
The pH dependence of the thermal stability. The temperature melting curves of CMPs at pH 3.6 (A), and pH 2.9 (B); the same color code in Fig. 2 was used for α1C (blue), α1C-K(cyan), α2C (orange), α2CP (red), α2CP-K (light magenta); the blue dashed curves in A and B are the melting curve of α1C at pH 7.0 included for comparison. The colored arrows in B are used to highlight the decreased *T_m_* of α1C-K and α2CP comparing to α1C (cyan and red, respectively), and α2CP-K comparing to α2CP (magenta). C) the temperature melting curves of peptide α2N in pH 2.9 (diamond), pH 3.6 (triangle), pH 7 (circle), and pH 9.4 (square). D) the amino acid sequences of peptides at pH 2.9 showing the positively charged residues in blue, and protonated Glu and Asp in light pink.

Although protonation of the His-tag lowered the *T_m_* of all peptides by ∼ 5 °C, it had little effect on their relative thermal stabilities, nor did it alter the approximately two-state character of the thermal transition. α1C and α2C-P remained the most stable constructs, whereas α2C remains the least stable one with a *T_m_* about another 3 °C lower than the others. Most interestingly, the His-tag induced unfolding at pH 3.6 did not alter the effects—or lack of thereof—of the inserted KGE sequences. The *T_m_* values of α1C and α1C-K remained nearly identical, so did those of α2C-P and α2C-PK (Fig. 3A and Table 1) despite partial protonation of Glu residues (pKa of 4.3). The persistent lower *T_m_* of α2C relative to the others at pH 3.6, on the other hand, suggests additional destabilizing effects which may have been exacerbated by the low pH and the N-terminal unfolding. The stabilizing effects of the Ile-to-Pro substitution in α2C-P is not sensitive to pH and contribute to the approximately ∼ 2 °C stability advantage relative to α2C (Fig. 3A and Table 1). The lowered pH, however, is expected to protonate the His-residues of the Gly–Lys–His tripeptide at position 883-885 of α2C (Fig. 1A) generating a cluster of six closely spaced positive charges across the three chains of the triple helix close to the N-terminus, which is a potentially highly unfavorable electrostatic configuration. The lower frequency of analogous motif such as Gly-Lys-Lys or Gly-Asp-Glu tripeptide in fibrillar collagens may reflect the structural penalty associated with concentrating like charges within a single tripeptide (21).

Lowering the pH to 2.9 causing the protonation of the Asp and Glu side chains in addition to His, revealed additional sequence dependent differences in thermal stability (Fig, 3B and Table 1). While the α1C showed minimal additional change in *T_m_* at pH 2.9, α2C-P exhibited a *T_m_* nearly ∼ 4 °C lower than α1C. Both peptides carry substantial net positive charge at this pH, with α2C-P having a net charge of approximately +10 and α1C approximately +9, although the contribution of individual charged residues is likely dependent on X- or Y-position and strongly modulated by the sequence context. A noticeable difference is an extra Y-position Arg in α1C which is known for its stabilizing effects because the longer Arg side chain can form favorable H-bonding interactions and van der Waals interaction with the peptide backbone (11, 21, 26). Regardless, the increased electrostatic repulsion at pH 2.9 appears sufficient to partially overcome the stabilizing effect of the Ile-to-Pro substitution in α2C-P. Without this Ile-to-Pro substitution, however, the triple helix is even less stable: α2C exhibited a *T_m_* of 23.4 °C, ∼ 3 °C lower than α2C-P, and more than 7 °C lower than α1C. The two peptides with the KGE insertions were further destabilized at pH 2.9 each showing an approximately 2 °C reduction in *T_m_* compared to their parent peptides which retained the original Y-position Arg. At this pH, the proposed interchain salt bridges are no longer possible because the Glu side chains are fully protonated. Instead, the reduced stability is consistent with the stabilizing effects of Y-Arg for the reason mentioned above. Yet again, these destabilizing interactions only shifted *T_m_* values while preserving the apparently cooperative nature of the thermal transition.

Despite having an identical triple-helical sequence as α2C, α2N is substantially more stable than α2C (Fig. 3C, Table 1) because the N-terminal Cys-knot in α2N not only suppresses His-tag-induced unraveling but also constrains the conformational space accessible to the three chains in the unfolded state, thereby reducing the entropic cost of refolding. Unlike conventional crosslinking of free collagen peptides, whose stabilizing effect largely arises from eliminating the nucleation step, the foldon domain ensures that nucleation remains intact. Consequently, the higher stability of α2N directly reflects stabilization of the folded triple helix itself. Compared with α2C at neutral pH, the markedly higher Tₘ of α2N indicates that the same triple-helical sequence can exhibit markedly different thermal behavior depending on how unfolding is initiated. Thus, the thermal behavior of a region within a collagen triple helix is determined not only by its intrinsic sequence-dependent stability but also by how its unfolding is energetically coupled to neighboring regions. The small decrease in *Tₘ* observed at low pH may therefore reflect the charge effects on the intrinsic stability of the triple-helix arising from disruption of interchain electrostatic interactions from protonation of charged residues and the weakened H-bonds within the triple-helical domain itself. In contrast, the drastically reduced thermal stability of α2C at low pH, and of α2C-P for that matter, appear instead to result from the same charge-induced destabilization that becomes amplified once unfolding is initiated at the N-terminus.

The N-terminal unfolding observed here therefore provides an experimental model of local unfolding initiation. Although protonation of the His-tag produces a similar downward shift in melting temperature for all constructs without N-terminal crosslinking, the persistence of distinct melting temperatures demonstrates that intrinsic sequence-dependent differences remain. Taken together, these findings suggest that unfolding is preferentially initiated in regions of lower intrinsic stability, whereas the continuity of the triple helix governs its subsequent propagation. Whether this influence persists as the helix becomes substantially longer is examined below.

### The thermal transition of extended triple helices

If local unfolding initiation governs the apparent thermal transition of the triple-helix, increasing the distance between that initiation site and the remainder of the helix might be expected to reduce its influence. To test this prediction, we expanded the N-terminal GPP_4_ segment of α1C to (Gly-Pro-Pro)_6_, adding two more stabilizing GPP triplets. The additional (Gly-Pro-Pro) tripeptides increased the *T_m_* of α1C-GPP_6_ by more than 3 °C to *∼* 41.08 °C (Fig. 4A, Table 1), but did not reduce the susceptibility of the helix to unfolding at the N-terminus. α1C-GPP_6_ displayed similar pH dependence, with minimal change at pH 9.4 but a substantial ∼ 3 °C decrease at pH 3.6. These effects were reproducible in both the presence and absence of added salt (Fig. 4A). Thus, even after increasing the intrinsic stability and increasing the distance from the perturbation, the helix remains highly susceptible to the unfolding initiation.

**Figure 4.**
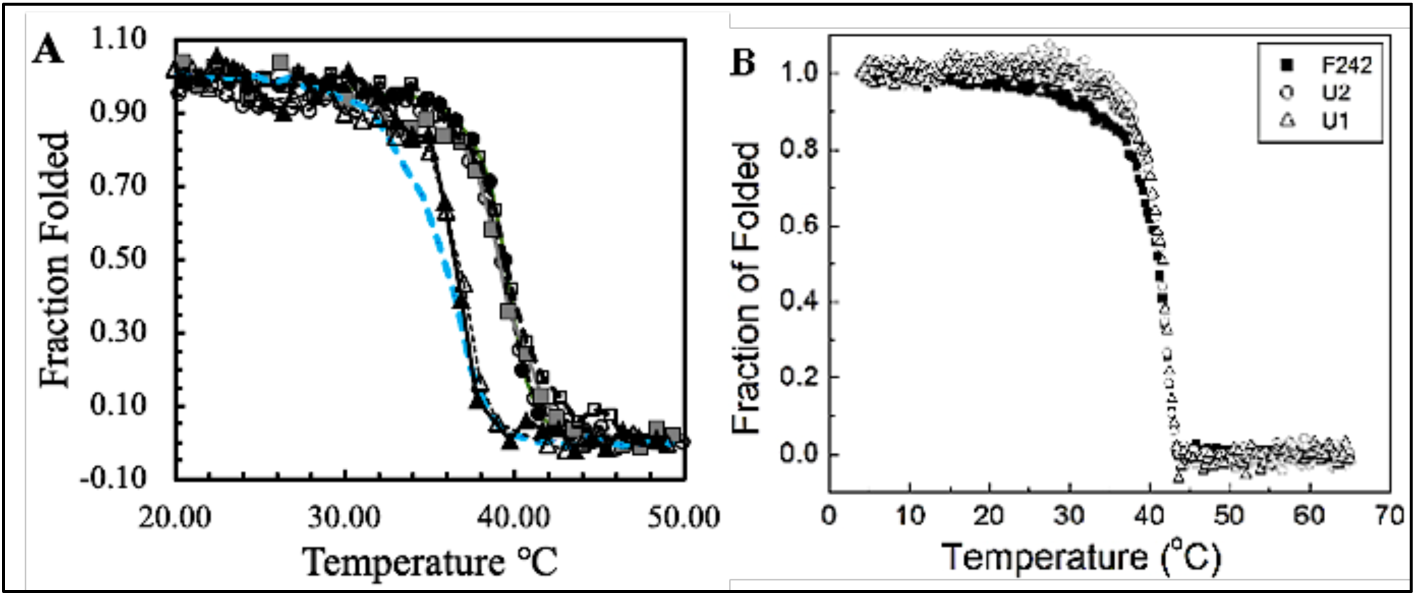
the dependence of the triple helix stability on the length of the peptides. A) the temperature melting curves of peptide Trix^-^ in buffers of pH 3.6 (triangle), pH 7.0 (circle), and pH 9.4 (square) without salt (solid lines and filled symbols) or with salt (dashed lines and open symbols); the melting curve of α1C at pH 7.0 is also included for comparison (blue dash). B) the temperature melting curves of peptides having different length of the triple helix domain: U1 (128 residues), U2 (267 residues), and F242 (360 residues), in pH 7.0 PBS buffer.

Interestingly, the melting temperature of α1C-GPP_6_ (41.08 °C) is remarkably close to that of another peptide developed in our lab, Col-877, which exhibits a *T_m_* of ∼ 42 °C despite being nearly three times longer (27). Col-877 consists of three tandem repeats of a sequence unit compromising the same Col-domain found in α1C and α1C-GPP_6_ together with a GPP_4_ segment. Addition of two GPP triplets increased the *T_m_* by more than 3.5 °C, while tripling the overall size of the triple-helix produced only an additional ∼1 °C increase in thermal stability. Clearly, the observed thermal transition is not simply the cumulative consequences of local stabilizing interactions, despite the strong sequence-dependence of intrinsic stability.

To determine whether this behavior is unique to the α1C sequence or represents a more general property of extended collagen triple helices, we examined a second series of rCMPs. Similarly, no detectable differences in *T_m_* were found among peptides containing triple-helical domains of approximately, 138, 224, and 352 residues (Fig. 4B, (27, 28)). The shortest peptide in this series, U1, is structurally similar to α1C-GPP_6_ but contains a longer Col-domain (108 vs 84 residues) with a different amino acid sequence (28, 29). Peptides U2 and F242 contains two and three tandem repeats, respectively, of the U1 sequence unit, and all three peptides possess an N-terminal GPP_6_ segment analogous to that in α1C-GPP_6_. Other studies from our laboratory of rCMPs with triple helix domains ranging from ∼ 100 to more than 300 residues also indicate that the triple helices exhibit remarkably similar thermal transition despite substantial sequence variations, helix length, and in some cases, the use of different nucleation domains (27–31). Because nucleation is maintained in these rCMPs at the C-terminus by the foldon domain and Cys-knot, thermal unfolding is therefore initiated predominantly from the opposite end. In addition to the negligible changes in *T_m_* values, the apparent two-state thermal transitions were also uniformly observed. Thus, the weak dependence of the observed thermal transition on chain length suggests that the effective unfolding unit of extended helices is not determined by the size of individual stability domains but the continuity of the triple-helix itself.

The diminishing dependence of *Tₘ* on chain length is readily explained by the increasing entropic cost of folding extended chains, which progressively offsets the additional stabilizing interactions contributed by longer triple helices. This enthalpy–entropy compensation explains why *Tₘ* eventually reaches diminishing returns with increasing chain length, but it does not explain why the unfolding transition itself remains highly cooperative. Our observations instead suggest that this behavior may arise from the continuity of the triple helix itself: once unfolding is initiated within a locally unstable region, the folded neighboring regions remain structurally coupled, allowing unfolding to propagate cooperatively through the entire helix. Such cooperative propagation would effectively couple regions of differing intrinsic stability into a single unfolding event, thereby masking their individual thermodynamic transitions. This interpretation is also consistent with previous studies of bacterial collagens and human type II collagen, in which regions of differing intrinsic thermal stability could be identified in isolated fragments, whereas the intact molecules nevertheless exhibited a single, approximately two-state thermal transition (ref). This interpretation reconciles the apparent discrepancy between the identification of local stability domains in isolated collagen fragments and the predominantly two-state thermal transitions observed for intact collagen molecules, providing a mechanistic framework for understanding how intrinsic thermodynamic heterogeneity is integrated within extended collagen triple helices.

## CONCLUSIONS

This study demonstrates that the intrinsic thermal stability of the collagen triple helix remains strongly sequence dependent despite the apparently two-state character of its thermal transition. Individual stabilizing interactions from Pro and Y-Arg remain largely preserved in extended triple helices, whereas the interchain salt bridges form KGE depend strongly on their surrounding sequence context, indicating that local sequence continues to govern the thermodynamics of the folded state. At the same time, the pronounced destabilization caused by N-terminal unfolding shows that identical triple-helical sequences can exhibit markedly different thermal behavior depending on how unfolding is initiated.

The persistence of an approximately two-state thermal transition across peptides differing substantially in sequence and length further indicates that the observed thermal transition of an extended collagen triple helix is governed by more than the cumulative sum of local stabilizing interactions. Instead, our findings support a mechanism in which unfolding is preferentially initiated within regions of lower intrinsic stability, whereas the continuity of the triple helix thermodynamically couples neighboring regions into a cooperative unfolding process. Such coupling provides a plausible explanation for the longstanding paradox that intact collagen molecules generally exhibit a single thermal transition despite accumulating evidence for regions of differing intrinsic stability.

These findings further suggest that residue propensities derived from host–guest peptides should be interpreted primarily as measures of intrinsic local thermodynamic stability rather than as direct predictors of the behavior of individual regions within an extended collagen triple helix. Regions of lower intrinsic stability are therefore not expected to behave as independently unfolding domains, because their unfolding remains coupled to neighboring regions of the continuous triple helix. Likewise, intrinsically stable regions may nevertheless unfold once cooperative propagation has been initiated elsewhere. The biological significance of local thermodynamic heterogeneity may therefore lie less in defining independently unfolding domains than in modulating how individual regions respond to perturbations within the context of the continuous triple helix.

## ASSOCIATED CONTENT

The Supporting Information includes SDS-PAGE analysis of purified recombinant collagen mimetic peptides and representative thermal denaturation experiments demonstrating reproducibility.

## AUTHOR CONTRIBUTIONS

S.Y.X. performed most of the experiments and analyzed the data. S.W. created the peptide series used to study sequence and length dependence of extended triple helices, performed the experiments and analyzed the data. S.T. designed α2C and α2N peptides and performed the experiments. F.A. participated in the purification and characterization of α2N peptide. Y.X. conceived and supervised the study, analyzed and interpreted the data, and wrote the manuscript. All authors reviewed and approved the final manuscript.

## FUNDING SOURCES

This work was supported in part by NSF grant CHE 1022120, NIH grant SC1 GM121273, and PSC-CUNY Research Award Programs (Grant No. 6871156 A, and ENHC 6668954)

## ACKNOWLEDGMENTS

The authors thank Drs. Anton Persikov and Nathan Astrof for insightful discussions and critical reading of the manuscript.

## Notes

### Competing Interest Statement

The authors have declared no competing interest.

