## Supplemental Table S1, Fig. S1, and Fig S2 for "Sequence-dependent Stability and the Apparent Two-state Thermal Transition of Extended Collagen Triple Helices"

Contents

- Figure S1. SDS-PAGE analysis of purified rCMP α1C ............ S2
- Table S1. The molecular weight and properties of the rCMPs ............ S3
- Figure S2. Reproducibility of representative thermal denaturation experiments ............ S4

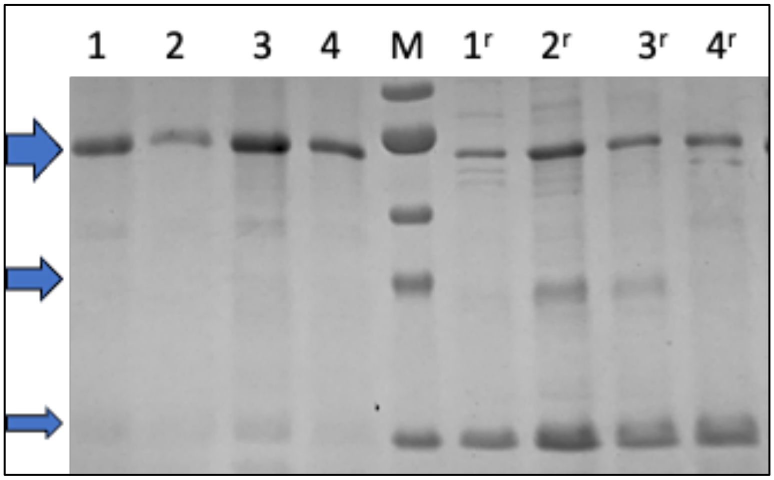


**Figure S1. SDS-PAGE analysis of purified rCMP α1C.** Lanes 1 and 2, preparation 1 at 0.40 and 0.18 mg/mL, respectively; lanes 3 and 4, preparation 2 at 0.319 and 0.150 mg/mL, respectively. Lanes 1r–4r correspond to lanes 1–4 after reduction with 0.36 M DTT. Lane M, molecular weight markers (65, 52, 37, 30, and 16 kDa). Blue arrows indicate the positions of the trimer (≈39 kDa), dimer (≈26 kDa), and monomer (≈13 kDa). Owing to their elongated triple-helical structure and amino acid composition, collagen triple helices typically exhibit reduced electrophoretic mobility and therefore migrate at an apparent molecular weight higher than expected from their calculated molecular mass. Gel concentration: 15%.

**Table S1. Molecular weight and peptide length of the rCMPs**

| rCMP | Molecular weight (Da) | Amino acid length | Theoretical PI | Total negatively charged amino acids | Total positively charged amino acids |
|---|---|---|---|---|---|
| α1C-$GPP_6$ | 13468.11 | 139 | 8.76 | 10 | 12 |
| α1C | 12930.26 | 133 | 8.76 | 10 | 12 |
| α1C-K | 12902.24 | 133 | 8.73 | 10 | 12 |
| α2C | 13045.52 | 133 | 9.39 | 8 | 12 |
| α2P | 13029.48 | 133 | 9.39 | 8 | 12 |
| α2P-K | 13001.47 | 133 | 9.34 | 8 | 12 |
| α2N | 13045.52 | 133 | 9.39 | 8 | 12 |

All constructs' single-chain peptide molecular weights and amino acid lengths are listed. All values are calculated using the ProtParam tool using the sequencing data.

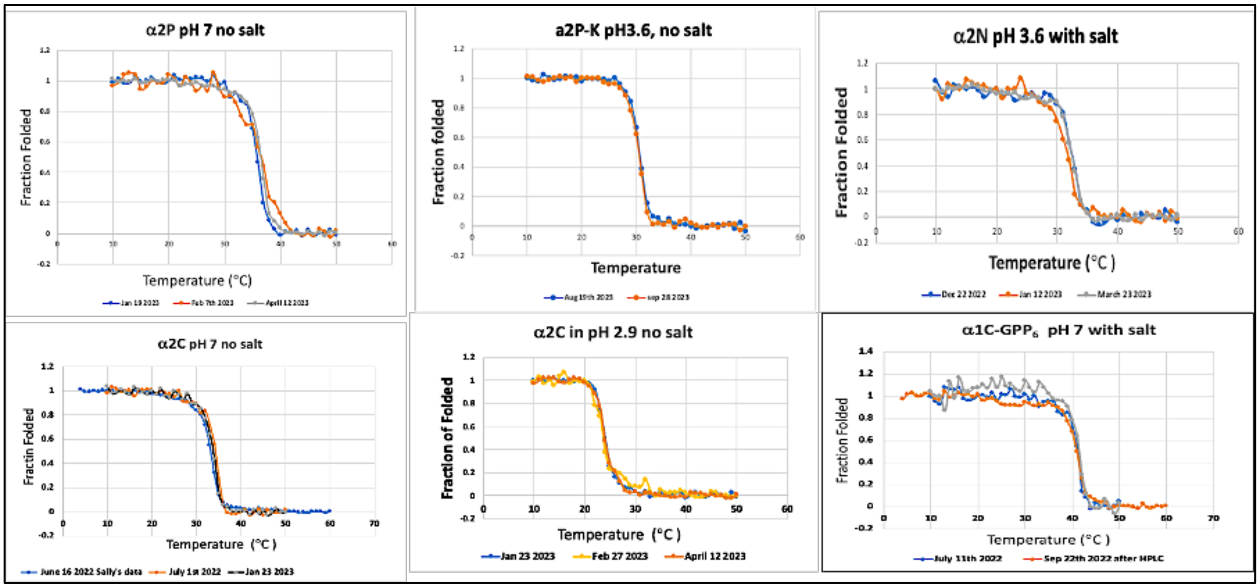


**Figure S2. Reproducibility of representative thermal denaturation experiments.** Thermal denaturation curves of each recombinant collagen mimetic peptide (rCMP) obtained from independent protein preparations at different peptide concentrations under identical buffer conditions and a heating rate of 1 °C/min. The similar transition temperatures observed at different peptide concentrations demonstrate that thermal unfolding is independent of peptide concentration, consistent with stabilization of the triple helix by the C-terminal foldon domain and interchain disulfide crosslinking.
